# Pan-cancer analysis of mRNA stability for decoding tumour post-transcriptional programs

**DOI:** 10.1101/2020.12.30.424872

**Authors:** Gabrielle Perron, Pouria Jandaghi, Maryam Rajaee, Rached Alkallas, Yasser Riazalhosseini, Hamed S. Najafabadi

## Abstract

RNA stability is a crucial and often overlooked determinant of gene expression. Some of the regulators of mRNA stability are long known as key oncogenic or tumour suppressor factors. Nonetheless, the extent to which mRNA stability contributes to transcriptome remodeling in cancer is unknown, and the factors that modulate mRNA stability during cancer development and progression are largely uncharacterized. Here, by decoupling transcriptional and post-transcriptional effects in RNA-seq data of 7760 samples from 18 cancer types, we present a pan-cancer view of the mRNA stability changes that accompany tumour development and progression. We show that thousands of genes are dysregulated at the mRNA stability level, and identify the potential factors that drive these changes, including >80 RNA-binding proteins (RBPs) and microRNAs (miRNAs). Most RBPs and miRNAs have cancer type-specific activities, but a few show recurrent inactivation across multiple cancers, including the RBFOX family of RBPs and miR-29. Analysis of cell lines with phenotypic activation or inhibition of RBFOX1 and miR-29 confirms their role in modulation of genes that are dysregulated across multiple cancers, with functions in calcium signaling, extracellular matrix organization, and stemness. Overall, our study highlights the critical role of mRNA stability in shaping the tumour transcriptome, with recurrent post-transcriptional changes that are ~30% as frequent as transcriptional events. These results provide a resource for systematic interrogation of cancer-associated stability drivers and pathways.

## Background

Widespread disruption of gene expression programs is a hallmark of cancer and underlies the extensive transformation of tumour cell identity and behavior. Among the least understood aspects of this gene expression remodeling is the regulation of mRNA stability and decay. Previous studies have found specific programs that are involved in tumorigenesis or metastasis through modulation of mRNA stability [1–8]; however, the extent to which mRNA stability contributes to cancer cell transcriptome has not been systematically studied, and the associated regulatory networks are mostly unknown. A key limitation in studying these post-transcriptional programs stems simply from our lack of ability to measure mRNA decay rate *in vivo*: traditional methods that measure mRNA decay rely on *in vitro* manipulations such as transcriptional inhibition with chemical inhibitors (e.g. actinomycin D) or metabolic labeling with nucleoside analogues (e.g. 4-thiouridine), combined with time series measurements of transcripts [9–11]. Despite recent improvements [12, 13], these methods are resource-intensive, have inherent limitations and biases such as triggering cellular stress and pleiotropic effects [14], and, most importantly, are only applicable to *in vitro* models. As a result, the mRNA stability landscape of tumour remains almost completely uncharted across different cancer types.

A potential solution comes from recent studies showing that tissue RNA-seq data contain enough information to disentangle transcription rate from mRNA decay rate. Briefly, under the assumption that RNA processing rate is constant [15, 16], any change in unspliced (pre-mature) mRNA abundance (estimated from intronic reads) must reflect a proportional change in transcription rate, while any change in spliced (mature) mRNA abundance (estimated from exonic reads) reflects the combined effect of transcription rate and mRNA decay (Fig. 1a). This model enables the estimation of differential mRNA stability based on how the ratio of exonic and intronic reads changes across conditions [15]. A recent improvement on this model generalizes the unspliced-spliced relationship as a power-law function, with the power-law exponent reflecting the coupling between transcription rate and splicing rate [17] (**Fig. S1a,b**).

**Figure 1.**
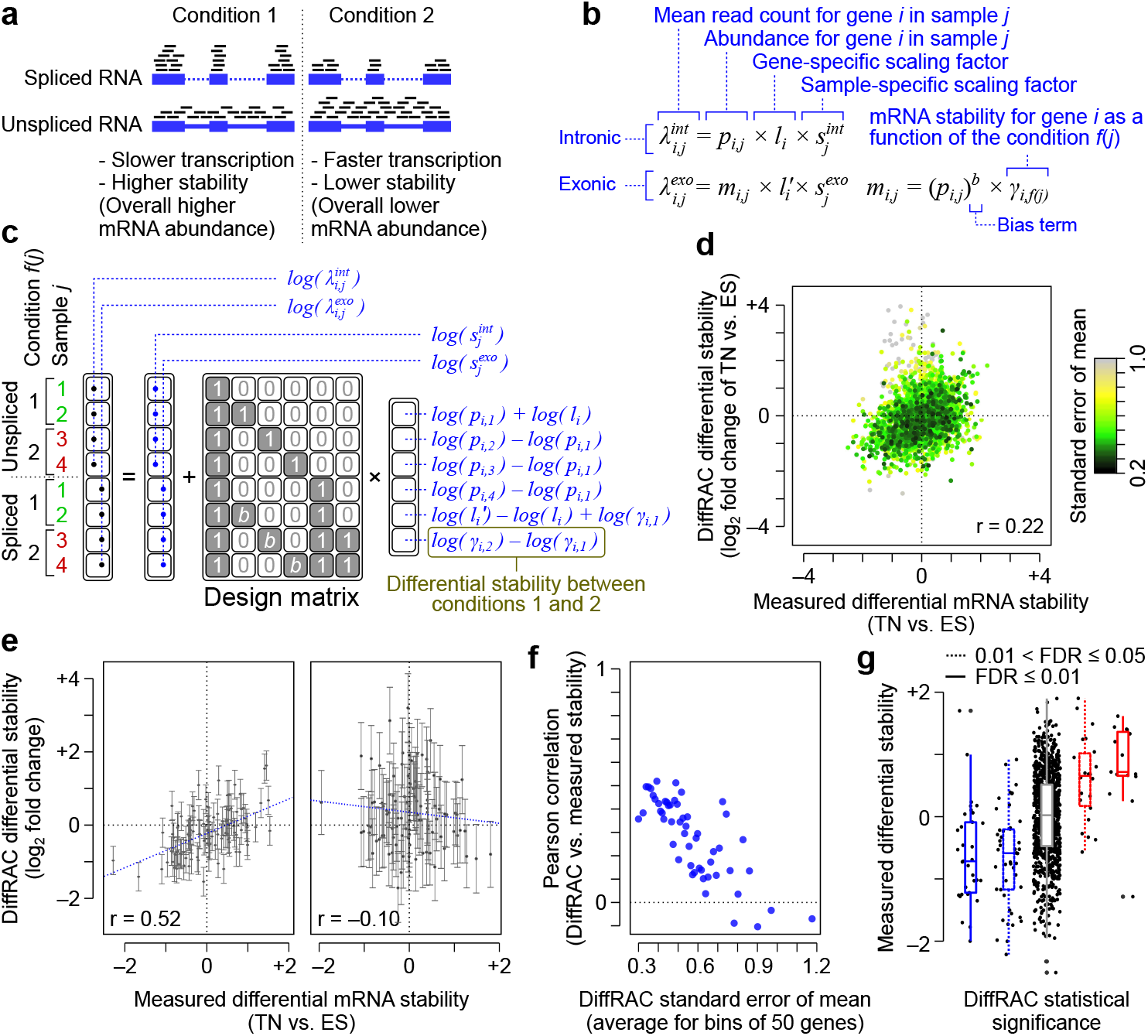
Inference of differential mRNA stability using DiffRAC. (**a**) Schematic representation of the effect of transcription and stability on the abundances of unspliced and spliced RNA. (**b**) DiffRAC models the mean (*λ*) of intronic (*int*) and exonic (*exo*) read distribution as a function of pre-mature (*p*) and mature (*m*) transcript abundances, in addition to gene-specific (*l*) and library-specific (*s*) scaling factors. The mature mRNA abundance is itself modeled as a function of the pre-mature RNA abundance and mRNA stability (*γ*), which is in turn a function (*f*) of the experimental variables. Also see Fig. S1. (**c**) An example case with four samples and two experimental conditions, showing how DiffRAC’s model can be implemented in a regression with a log-link function, along with the interpretation of regression coefficients (also see Methods). (**d**) Comparison of DiffRAC stability estimates against experimental mRNA half-life (stability) measurements in mouse ES cells differentiated to terminal neurons (TN) [18]. Each data point stands for one gene, with the points colored according the standard error of the mean (SEM) for DiffRAC estimates. (**e**) Comparison of DiffRAC estimates vs. measured stability for the 100 genes with the smallest (left) and largest (right) DiffRAC SEMs. Error bars represent SEM. (**f**) The Pearson correlation between DiffRAC estimates and measured stability for bins of 50 genes sorted by their SEM. (**g**) Distribution of experimental half-life measurements for genes that DiffRAC has identified as significantly destabilized (blue boxplots) or stabilized (red boxplots) in TN vs. ES cells, at FDR cutoffs of 0.05 (dashed line) or 0.01 (solid line). Genes that are not called as significant by DiffRAC are represented with the grey boxplot.

Here, we build on these methods to obtain a pan-cancer map of mRNA stability changes between tumour and normal tissues, as well as the mRNA stability changes that accompany tumour progression. To do so, we first introduce a general framework for statistical analysis of differential mRNA stability that takes into account the distributional properties of count data. We benchmark this method using experimental measurements of mRNA decay rate, and then apply it to the RNA-seq data from The Cancer Genome Atlas (TCGA) to map the mRNA stability landscapes of 18 cancer types. We identify thousands of transcripts whose stability is altered during tumour formation and/or progression – experimental measurements in cancer cell line models support these findings and suggest a role for mRNA stability alterations in tumor progression and invasiveness. Finally, using network modeling and functional experiments, we identify key microRNAs (miRNAs) and RNA-binding proteins (RBPs) that mediate these changes, providing new insights into the post-transcriptional mechanisms of transcriptome remodelling in cancer.

## Results

### A generalized linear model for statistical testing of mRNA stability

The spliced and unspliced transcripts of each gene follow a power-law relationship, with deviations from this power-law trend reflecting changes in the degradation rate of the mature mRNA [17] (**Fig. S1a,b**). The power-law exponent reflects the coupling between transcription rate and RNA processing rate – an exponent of 1 indicates no coupling between transcription and processing rate constants, whereas values smaller than 1 indicate that as transcription increases, processing rate constant decreases, potentially due to saturation of the RNA processing machinery (**Fig. S1a**). To use this power-law relationship for the inference of differential stability, it is essential to correctly model the variability in RNA-seq counts. For this purpose, we developed DiffRAC (https://github.com/csglab/DiffRAC), a framework that converts the unspliced-spliced relationship to a generalized linear model whose parameters can then be inferred from sequencing count data using an appropriate error model of choice (**Fig. 1b-c** and **Fig. S1c-d**).

We evaluated the performance of DiffRAC for estimating differential mRNA stability using a previously published benchmarking dataset [18], consisting of RNA-seq data from mouse embryonic stem cells and terminal neurons, along with ground-truth transcript half-life measurements after transcriptional blockage with actinomycin D. We observed a modest overall correlation between RNA-seq-based stability estimates from DiffRAC and ground-truth stability measurements (Pearson correlation 0.22; **Fig. 1d** and **Table S1**), in line with previous reports on RNA stability estimation using this specific benchmarking dataset [17, 18]. However, for genes that had narrow confidence intervals as estimated by DiffRAC, the Pearson correlation between RNA-seq-based estimates and ground truth exceeded 0.5 (**Fig. 1d-f**), indicating that the confidence intervals estimated by DiffRAC indeed reflect the true uncertainty in estimating differential mRNA stability. Based on (adjusted) P-values associated with DiffRAC differential stability estimates, we identified 79 genes with higher stability in embryonic stem cells and 37 genes with higher stability in terminally differentiated neurons (FDR < 0.05), which closely correspond to differentially stable genes based on the ground truth (**Fig. 1g**). Overall, these results suggest that DiffRAC can properly estimate not just the mean differential mRNA stability, but also its uncertainty and statistical significance.

One limitation of the model described above is that, with increasing sample sizes, the number of latent variables that need to be estimated by regression also increases, which can become prohibitively expensive in terms of computational times. To overcome the challenges associated with fitting the model in large sample cohorts, we developed a simplified DiffRAC model that assumes most of the variance in transcription can be explained by the experimental variables (see **Methods** and **Fig. S2a-c**). This assumption greatly reduces the number of parameters; however, we observed that it does not significantly alter the differential stability estimates in the benchmarking dataset (**Fig. S2d**).

### DiffRAC identifies cancer-associated changes in mRNA stability

To investigate the post-transcriptional changes responsible for transcriptome remodeling in cancer, we performed a pan-cancer analysis of differential mRNA stability across TCGA, encompassing 7760 samples from 18 cancer types. We used DiffRAC to identify genes that were differentially stabilized or destabilized in tumour compared to normal tissues in each cancer type. This analysis revealed an average of 3954 genes that were differentially stabilized/destabilized per cancer type (FDR-adjusted p < 0.05) (**Fig. 2a-b**, **Fig. S3**, and **Table S2**), suggesting widespread post-transcriptional remodeling in cancer. The majority of genes showed highly cancer-specific stability profiles (**Fig. 2b**).

**Figure 2.**
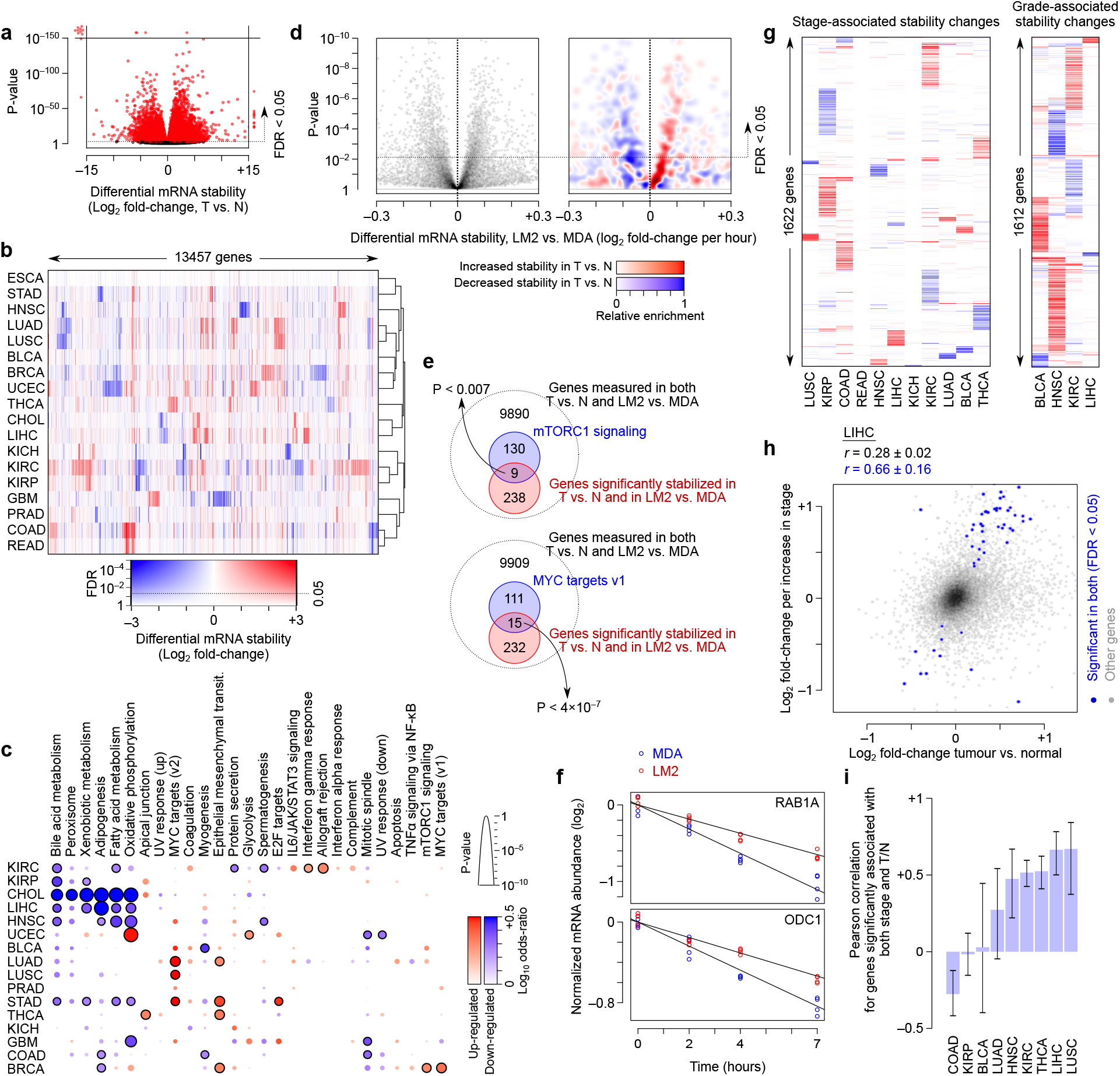
Pan-cancer analysis of differential mRNA stability. (**a**) Volcano plot of differential RNA stability between tumour and normal tissues for 18 TCGA cancer types. See Fig. S3 for volcano plots of individual cancer types. (**b**) Heatmap of differential mRNA stability profiles across TCGA cancers. Genes with significant DiffRAC results in at least one cancer (FDR < 0.05) are included. The color gradient represents a combination of the log2 fold-change of mRNA stability and the FDR. BLCA: bladder urothelial carcinoma, BRCA: breast invasive carcinoma, CHOL: cholangiocarcinoma, COAD: colon adenocarcinoma, ESCA: esophageal carcinoma, GBM: glioblastoma multiforme, HNSC: head and neck squamous cell carcinoma, KICH: kidney chromophobe, KIRC: kidney renal clear cell carcinoma, KIRP: kidney renal papillary cell carcinoma, LIHC: liver hepatocellular carcinoma, LUAD: lung adenocarcinoma, LUSC: lung squamous cell carcinoma, PRAD: prostate adenocarcinoma, READ: rectum adenocarcinoma, STAD: stomach adenocarcinoma, THCA: thyroid carcinoma, UCEC: uterine corpus endometrial carcinoma. (**c**) Pathway enrichment analysis of genes with significant differential mRNA stability in each cancer type. Circles with black outline correspond to MSigDB hallmark gene sets that are significantly enriched among cancer-stabilized (red) and cancer-destabilized (blue) genes (FDR < 0.05, Fisher’s exact test). Log-odds and P-values are represented using the color gradient and circle sizes, respectively. (**d**) The volcano plot on the left shows the experimentally measured differential stability between highly metastatic MDA-LM2 cell line relative to its parental MDA-MB-231 line (see Methods). The right panel shows the relative enrichment of genes that were stabilized (red) or destabilized (blue) in TCGA-BRCA tumors compared to normal samples. Kernel density estimation was used to calculate the density of BRCA-stabilized and destabilized genes across the plot, with the difference between the estimated densities of the two groups shown using the color gradient. (**e**) Venn diagrams illustrating the overlap between genes that are significantly stabilized in BRCA tumors (relative to normal) and MDA-LM2 (relative to MDA-MB-231), and genes that are part of the mTORC1 signalling (top) or MYC targets (bottom). P-values are based on Fisher’s exact test. (**f**) Transcription inhibition time course graphs for two example genes, one involved in mTORC1 signaling (RAB1A) and one among MYC targets (ODC1). Time course measurements in MDA-LM2 and parental MDA-MB-231 cells are shown in red and blue, respectively, with the slope of each fitted line representing the rate of degradation. (**g**) Stability changes associated with tumor stage (left) and grade (right) across TCGA cancers. Genes with significant changes in at least one cancer at FDR < 0.05 are included. The color gradient is the same as in panel **b**. (**h**) Comparison of the differential mRNA stability between tumour and normal (x-axis) and stage-associated differential mRNA stability (y-axis), in the TCGA-LIHC dataset as an example. Genes with significant changes along both axes at FDR < 0.05 are colored in blue. Pearson correlation coefficients and confidence intervals for all genes (black) and significant ones (blue) are shown on top. Panel (**i**) summarizes the Pearson correlations for significant genes in other cancer types (error bars represent the confidence intervals).

Several lines of evidence support the reliability of the stability profiles we have inferred. First, we observed that tumour mRNA stability profiles clustered by organ of origin (**Fig. 2b**), providing an internal validation for the robustness of stability inferences. Secondly, we observed that post-transcriptionally deregulated genes in each cancer type are functionally related (**Fig. 2c**), consistent with previously reported relationship between post-transcriptional regulons and functional gene modules [19, 20]. This analysis also highlights the role of mRNA stability in shaping the functional landscape of the cancer cell. For example, epithelial-mesenchymal transition genes and MYC targets are enriched among stabilized genes across several cancer types, while metabolic pathways such as oxidative phosphorylation and lipid metabolism are highly enriched among destabilized genes most noticeably in cholangiocarcinoma (CHOL), liver hepatocellular carcinoma (LIHC) and head-neck squamous cell carcinoma (HNSC).

Thirdly, we found that cancer-associated stability changes inferred from tissue RNA-seq data are highly consistent with experimentally measured mRNA stability changes in cancer cell line models. Specifically, we used time-series measurements of 4-thiouridine-labeled RNA [21] from MDA-MB-231 cell line, a model of breast cancer, as well as the highly invasive MDA-LM2 cells to identify mRNAs that are differentially stable between these two cell lines (see **Methods** for details; measurements are provided in **Table S3**). We found that the mRNAs that are more stable in breast tumours compared to normal tissue (based on DiffRAC analysis of tissue RNA-seq data) are also overall more stable in the highly invasive LM2 line compared to the parental MDA line (based on experimental stability measurements). Similarly, tumour-destabilized mRNAs that DiffRAC identified are overall less stable in the LM2 line (**Fig. 2d**).

Since the MDA-LM2 line is more invasive than MDA-MB-231, the above analysis also suggests that, at least in breast cancer, normal-to-tumour stability changes persist during the progression of the disease. In fact, two of the three pathways that were upregulated in breast tumors (**Fig. 2c**) also appear to be enriched among mRNAs that are stabilized in MDA-LM2 compared to MDA-MB-231 cell lines (MYC targets and mTORC1 signaling, **Fig 2e**; example genes are shown in **Fig. 2f**), supporting a role of mRNA stability in deregulation of these key pathways during both cancer development and progression. To understand whether normal-to-tumour stability changes are correlated with progression-associated stability changes across other cancers, we used DiffRAC to examine tumour stage and grade in each TCGA cancer type, while controlling for the confounding effects of age, sex, ancestry, and tumour purity (**Table S4**). We identified a total of 1966 genes with significant stability changes associated with tumor stage in at least one of the 11 cancers types that we analyzed (**Table S5**), and 2013 genes whose stability was associated with tumor grade in at least one of the four cancer types for which this type of classification was available (**Table S6**). We observed highly cancer-specific associations both for stage and grade (**Fig. 2g**). Importantly, we found that in most cases the stage- and grade-associated stability changes correlate with normal-to-tumour stability changes (**Fig. 2h** shows an example, with the overall results summarized in **Fig. 2i**). Taken together, these results highlight widespread mRNA stability changes in tumours, which affect key cancer-related pathways. This post-transcriptional deregulation continues to remodel the transcriptome through cancer progression, as evident from analysis of stage- and grade-associated stability changes.

### RNA-binding proteins play a key role in shaping the tumour mRNA stability profile

RNA-binding proteins (RBPs) and microRNAs (miRNAs) are the key regulators of mRNA stability. These sequence-specific factors primarily affect RNA stability through binding to the 3’ untranslated region (UTR) of their targets – RBPs either stabilize or destabilize their targets [22], while miRNAs primarily destabilize their target mRNAs [23, 24]. Starting with RBPs, we set out to examine whether these factors underlie the mRNA stability changes in cancer. We specifically tested for the enrichment of the targets of each RBP among genes that are differentially stable between tumour and normal tissues, after correcting for the background frequency of RBP binding to each gene (see **Methods**). **Figure 3a** shows an example, where the binding targets of the RBFOX1 protein are enriched among genes that are destabilized in glioblastoma multiforme (GBM), relative to the binding targets of other RBPs. We can quantify this enrichment by statistical modeling of the relationship between the binding of a specific RBP to the 3’ UTR of a gene and the tumour-specific stability status of that gene (**Fig. 3b**). We performed a systematic quantification of these relationships for 35 RBPs whose stability target sets (regulons) have been previously mapped based on the presence of their preferred binding sequences in the 3’ UTRs as well as the expression pattern of the candidate target genes [22]. This analysis revealed significantly enriched regulons among tumour-stabilized or destabilized genes across different cancer types, representing deregulation of 17 out of the 35 examined RBPs in at least one cancer type (**Fig. 3c**). Importantly, we observed excellent agreement between cancer-associated RBP expression changes and RBP target enrichments, after taking into account the expected function of each RBP in stabilizing or destabilizing its targets (Pearson correlation 0.61; **Fig 3d**). For example, SNRPA, which is an RNA-destabilizing factor [22], is up-regulated in multiple cancers, consistent with the observed destabilization of its regulon (**Fig. 3c-d**). This strong correlation highlights the reliability of our regulon analysis approach for identifying dysregulated RBPs, and suggests that aberrant expression of RBPs in cancer drives coordinated changes in the stability of their regulons.

**Figure 3:**
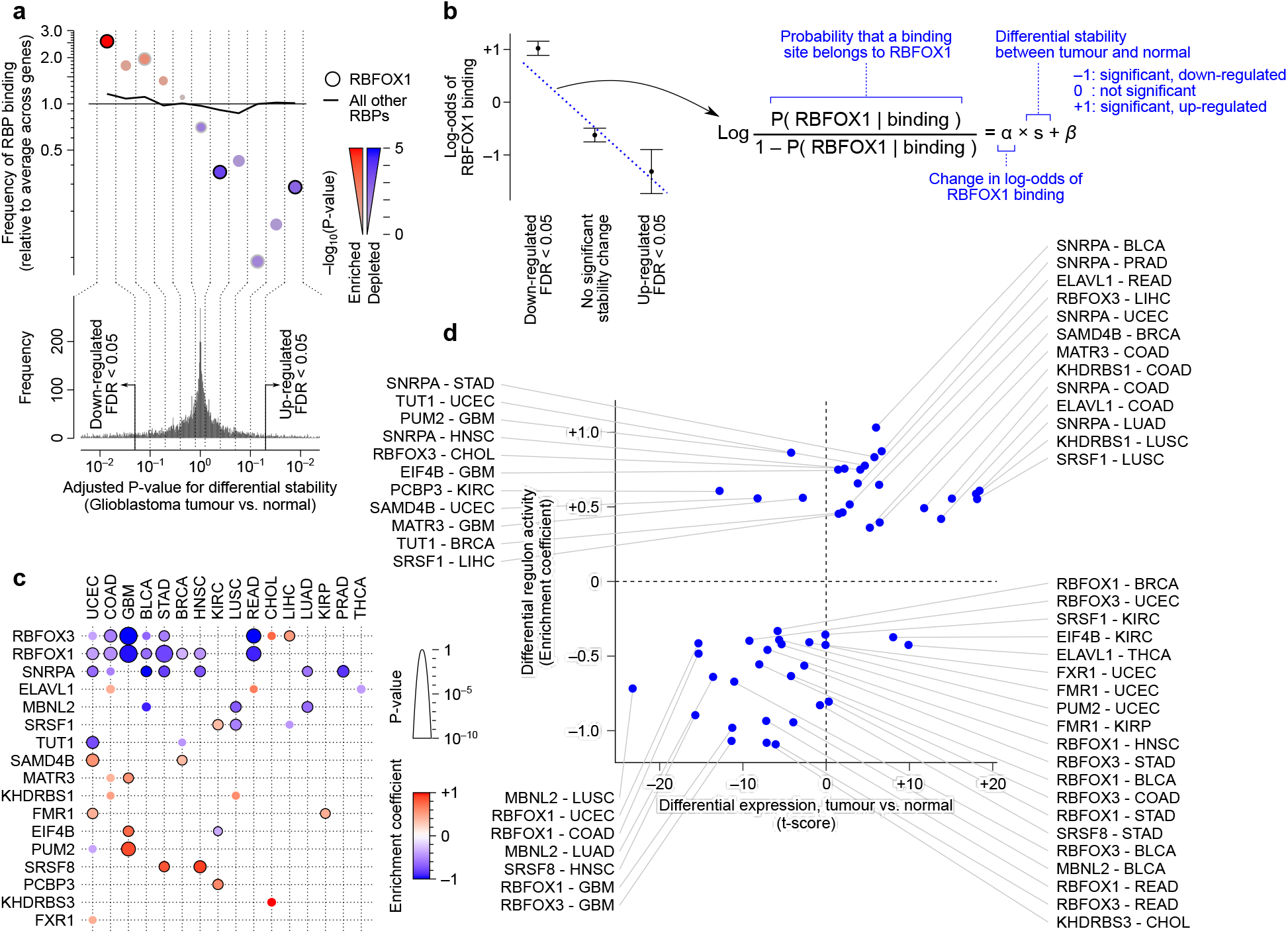
Enrichment of RBP binding sites among differentially stabilized genes in cancer. (**a**) An example case showing the enrichment of RBFOX1 binding sites among differentially stabilized genes in TCGA-GBM. Genes are binned by FDR of their DiffRAC differential stability between tumour and normal, with destabilized genes on the left and stabilized genes on the right. The relative frequency of RBFOX1 targets (circles) and targets of all other RBPs (solid line) is shown for each bin. (**b**) Schematic representation of the logistic regression approach for modeling the enrichment of RBFOX1 targets (relative to other RBPs) as a function of differential stability. (**c**) Heatmap summarizing the results of applying the model in panel **b** to all RBPs. Positive (red) and negative (blue) regression coefficients indicate enrichment of RBP targets among genes that are stabilized and destabilized in cancer, respectively. The circle size represents the significance level. Significant associations between RBP binding and stability status are shown using black outlines (FDR < 0.05). (**d**) Comparison of the differential RBP expression (tumor vs normal) and cancer-associated regulon activity. Regulon activity is defined to be the same as the enrichment coefficients from panel **c**, with the sign of the coefficient inverted for RBPs whose binding leads to RNA destabilization (based on [22]). Each dot represents one RBP in one cancer type. Pearson correlation of differential expression vs. differential regulon activity is 0.61.

Among the RBPs we analyzed, two RBPs, namely RBFOX1 and RBFOX3, stand out as being consistently deregulated across several cancer types. Specifically, the targets of these RBPs are enriched among destabilized genes in almost half of all the cancer types we analyzed (**Fig. 3c**). Consistent with the role of RBFOX proteins in promoting mRNA stability [22, 25], both RBFOX1 and RBFOX3 are down-regulated across multiple cancers (**Fig. 4a-b**), suggesting that down-regulation of RBFOX proteins leads to destabilization of their targets. For both RBFOX1 and RBFOX3, the highest expression in normal tissues can be seen in the brain tissue; subsequently, the most prominent case of their down-regulation as well as the most significant changes in the stability of their regulons can be seen in GBM, suggesting a major role in determining tumour transcriptome in this cancer type. However, their effect is not limited to GBM, especially for RBFOX3, which shows a broader range of expression in normal tissues and is also downregulated in a greater number of cancers (**Fig. 4b**).

**Figure 4:**
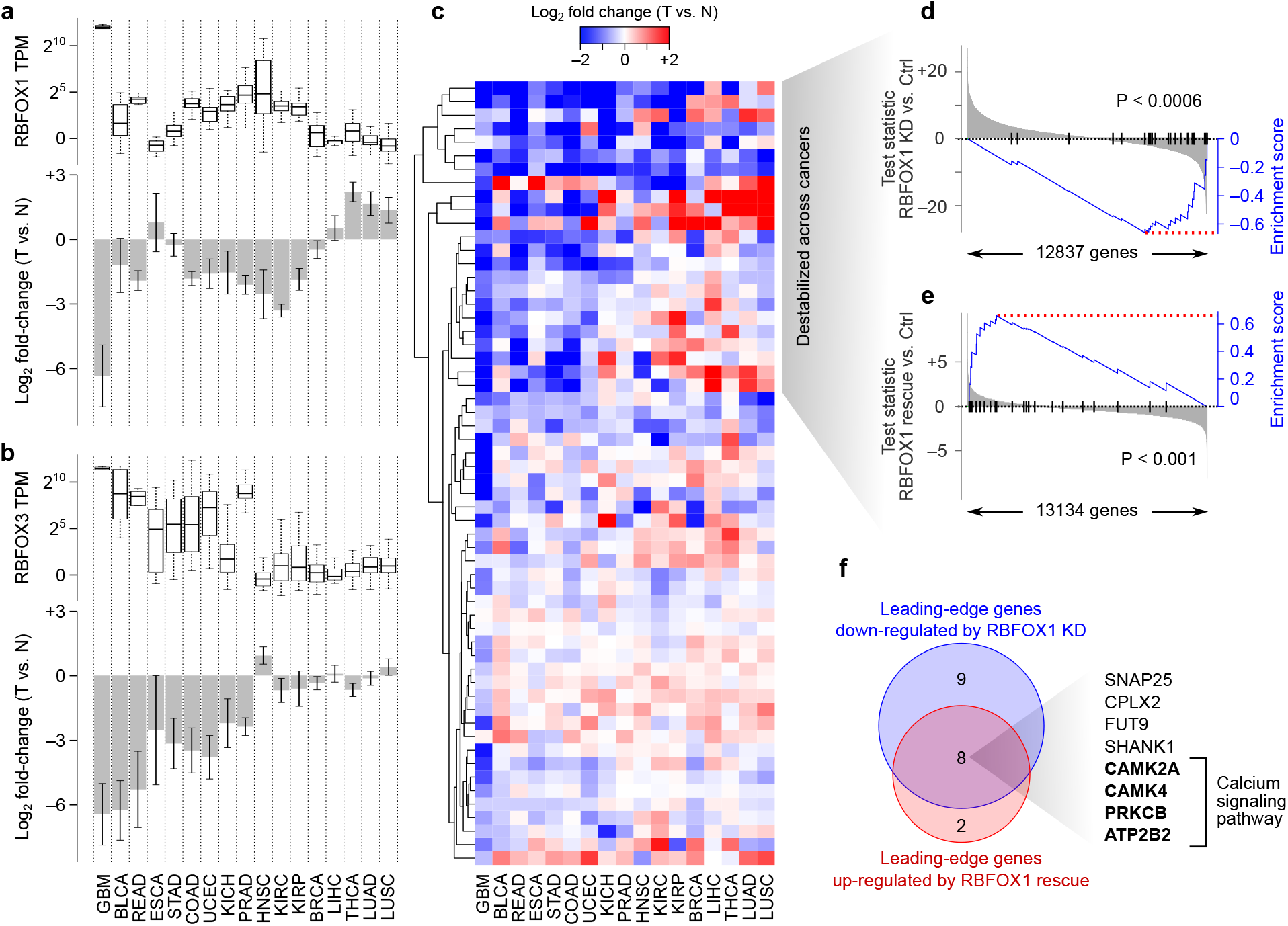
Aberrant activity of RBFOX proteins mediates stability changes across multiple cancers. (**a**) RBFOX1 expression across TCGA cancer types. The box plot (top) shows the RBFOX1 transcript-per-million (TPM) values in normal tissue samples. The bar plot (bottom) illustrates the average log fold-change of RBFOX1 expression in tumors compared to normal samples (error bars represent SEM). (**b**) RBFOX3 expression in normal tissue samples and differential expression in tumors, similar to panel **a**. (**c**) Heatmap showing the stability of RBFOX HITS-CLIP targets. HITS-CLIP targets include genes that have a mouse homolog with a strong RBFOX peak (height ≥200) in 3’ UTR, as defined previously [26]. Rows correspond to genes and columns to cancer types, with the latter sorted in the same order as panels **a**-**b**. (**d**) Gene set enrichment analysis (GSEA) [55] for RBFOX1 inhibition in terminally differentiated neurons. Genes (x-axis) are sorted by the Wald test statistic of differential expression between RBFOX1 knockdown and control cells, with vertical black lines demarcating the pan-cancer-destabilized set of RBFOX1 targets. The blue line represents the enrichment curve for this gene set [55]. (**e**) GSEA for RBFOX1 rescue in mouse neurons deficient for RBFOX proteins, similar to panel **d**. (**f**) Venn diagram illustrating the overlap between the leading-edge [55] set of genes downregulated by RBFOX1 knockdown (from **d**) and the leading-edge set of genes upregulated by RBFOX1 rescue (from **e**).

To confirm that the down-regulation of RBFOX proteins accompanies destabilization of their direct binding targets in cancer, we used HITS-CLIP data of RBFOX proteins in neurons [26] to build a high-confidence stability network, including genes that have the strongest binding sites in their 3’ UTRs (see **Methods**). As expected, we observed overall destabilization of HITS-CLIP-based RBFOX targets across different cancers (**Fig. 4c**). We noticed a subset of genes that are consistently destabilized across the same cancers in which either RBFOX1 or RBFOX3 is downregulated (**Fig 4c**). Interestingly, a subgroup of these genes is stabilized in the few cancer types in which RBFOX1 is upregulated (e.g. genes with positive stability values for LUSC, LUAD and THCA in **Fig 4c**), further supporting the notion that their cancer-associated stability changes are driven by RBFOX proteins.

To verify that the stability of these genes is regulated by RBFOX1, we examined the RNA-seq data from differentiated primary human neural progenitor (PHNP) cells in which RBFOX1 is knocked down [27]. As expected, cancer-destabilized genes that were associated with RBFOX1 were also downregulated upon RBFOX1 knockdown (**Table S7** and **Fig. 4d**). In contrast, when RBFOX1 expression is restored ectopically in the mouse neurons lacking RBFOX proteins [25], the expression of these genes is also rescued (**Fig. 4e**). We identified a core set of eight genes that have RBFOX binding site in their 3’ UTRs, are concurrently destabilized across cancers, are inhibited when RBFOX1 is knocked down, and are up-regulated when RBFOX1 expression is rescued (**Fig. 4f**). Interestingly, half of these genes belong to the calcium signaling pathway (based on KEGG pathways [28], Fisher’s exact test P<10^−6^), suggesting that deregulation of RBFOX proteins primarily affects calcium signaling in cancer cells.

### Dysregulation of miRNA regulons shapes the cancer transcriptome

To examine the contribution of miRNAs to the dysregulation of mRNA stability in cancer, we systematically searched for miRNAs whose targets are disproportionately dysregulated at the stability level in cancer, similar to the RBP analysis above (**Methods**). **Fig. 5a** shows miR-122 as an example; miR-122 is the most abundant miRNA expressed in liver cells [29], was previously shown to be downregulated in cholangiocarcinoma, and acts as a tumor suppressor via suppression of cell proliferation and induction of apoptosis [30, 31]. As expected, our regulon analysis indicates that miR-122 targets are predominantly stabilized specifically in cholangiocarcinoma tumours compared to normal tissue (**Fig. 5a**), consistent with reduced activity of miR-122. This observation is consistent with TCGA miRNA expression data, which show specific downregulation of miR-122 expression in cholangiocarcinoma (**Fig. S4**). Systematic application of this network-based approach revealed that, out of 153 broadly conserved miRNA families, the regulons of 63 miRNAs are deregulated in at least one cancer type, suggesting widespread disruption of miRNA networks (**Fig. 5b**).

**Figure 5:**
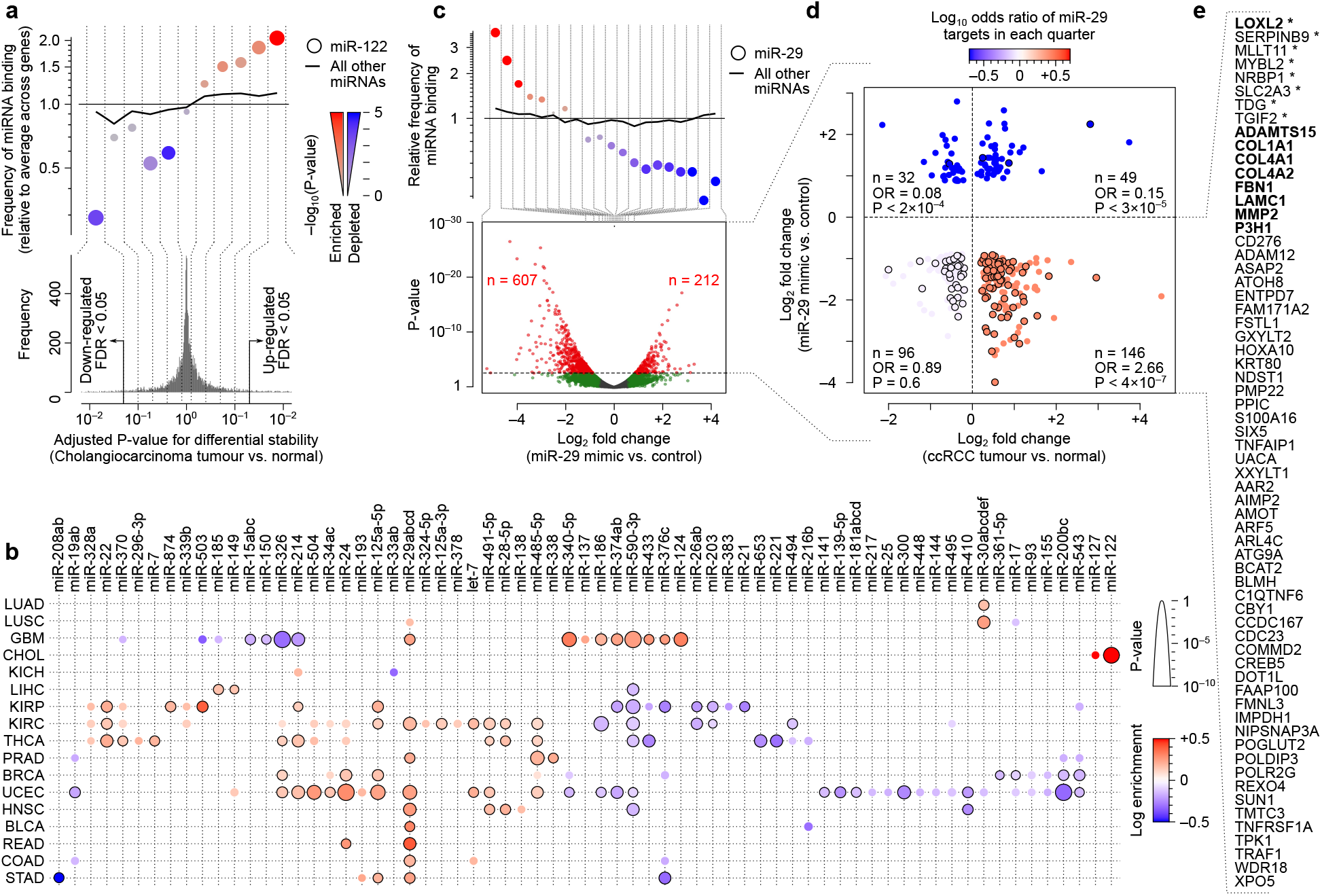
Dysregulation of miRNA regulons in cancer. (**a**) An example case showing the enrichment of miR-122 targets among genes stabilized in TCGA-CHOL tumours (relative to normal), similar to Fig. 3a. Note that since miRNAs are expected to destabilize their targets, enrichment in differentially stable genes indicates down-regulation of miRNA activity. (**b**) Heatmap summarizing enrichment analysis for all miRNAs across all cancer types, similar to Fig. **3c**. (**c**) Enrichment of miR-29 targets among genes that are destabilized after transfection of miR-29 mimic in 786-O cells. The volcano plot (bottom) summarizes differential expression results between miR-29 mimic and control; the dot plot at the top shows enrichment of miR-29 targets at bins of differentially expressed genes, similar to panel **a**. In the volcano plot, significantly differentially expressed genes (FDR < 0.05) are shown in red. (**d**) Enrichment of miR-29 binding sites, relative to other miRNA binding sites, in genes categories defined by their differential stability in TCGA-KIRC and differential expression after miR-29 mimic expression in 786-O cells. Each dot represents a gene, and those with a black outline contain at least one miR-29 binding site. The color gradient represents the log-odds of miR-29 binding site enrichment in each quarter. P-values are based on Fisher’s exact test. (**e**) List of genes that are bound by miR-29, up-regulated in KIRC, and down-regulated after miR-29-mimic treatment of 786-O cells. Genes that are bold correspond to ECM genes (based on overlap with GO; enrichment P < 6×10^−6^), and those with an asterisk are markers of embryonal carcinoma (based on StemCheker [56]; enrichment P < 3×10^−4^).

Of interest, we observed that miR-29 targets are recurrently stabilized across more than half of the cancer types we analyzed, suggesting a pan-cancer decrease in miR-29 activity. Among these cancer types, the miR-29 regulon showed the most significant enrichment among stabilized mRNAs in UCEC and KIRC (clear cell renal cell carcinoma), suggesting a major role in post-transcriptional remodeling in these cancer types. To understand whether restoring miR-29 activity can reverse these post-transcriptional changes, we expressed a miR-29 mimic in 786-O cells, which is a model for KIRC (**Fig. S5**). As expected, expression of miR-29 mimic resulted in global down-regulation of the miR-29 regulon (**Fig. 5c**). Importantly, miR-29 mimic expression leads to down-regulation of the majority of genes that are significantly stabilized in KIRC (**Fig. 5d**), most of which have a miR-29 binding site in their 3’ UTRs. Together, these results suggest that miR-29 down-regulation has a widespread effect on the stability of genes in cancer, while restoring its activity partially rescues the normal mRNA stability landscape of the cell.

## Discussion

By quantifying differential mRNA stability patterns across 18 cancer types, our study presents a systematic resource for mining the post-transcriptional landscape of the tumour. Importantly, our results uncovered recurrent changes in the stability of >13,000 genes in at least one cancer type, highlighting the widespread role of post-transcriptional regulation in shaping the cancer transcriptome. We note that this resource also provides an approximation for the relative contribution of transcriptional and post-transcriptional events in shaping cancer transcriptome: on average, 19% of genes that are significantly up-regulated at the expression level are detected by DiffRAC as significantly stabilized in tumour, and 23% of genes with significantly reduced expression are detected as significantly destabilized. In comparison, 66% and 61% of genes whose expression is significantly up- or down-regulated are detected as transcriptionally activated or inhibited in tumour, respectively (**Fig. S6**). These results suggest a significant role for post-transcriptional changes in shaping the cancer transcriptome, with recurrent changes that are ~30% as frequent as transcriptional events.

Our study also highlights the coordinated post-transcriptional deregulation of genes that are involved in the same pathways. Notably, we observed recurrent stabilization of mRNAs that encode epithelial-mesenchymal transition proteins and MYC targets across multiple cancer types. The epithelial-mesenchymal transition (EMT) is the process by which epithelial cells lose their apical-basal polarity and cell-cell adhesion, and instead acquire mesenchymal properties such as migratory and invasive potentials [32]; our results suggest that activation of the EMT pathway in cancer is at least partly mediated by post-transcriptional up-regulation. Similarly, we observed post-transcriptional up-regulation of MYC targets, which include growth-related genes that directly contribute to tumorigenesis [33]. MYC is a well-defined transcription factor and represents one of the most frequently amplified oncogenes [34], leading to transcriptional activation of its targets in cancer. Therefore, our intriguing observation that MYC targets are also up-regulated at the mRNA stability level suggests the presence of convergent transcriptional and post-transcriptional mechanisms that modulate overlapping gene sets. Furthermore, we observed coordinated destabilization of mRNAs for genes implicated in oxidative phosphorylation (OXPHOS) and related pathways such as fatty acid metabolism and adipogenesis, consistent with the well-documented Warburg effect in which upregulation of glucose consumption and glycolysis is accompanied by a down-regulation of OXPHOS [35].

In addition, we observed widespread and coordinated post-transcriptional modulation of the targets of RNA-binding proteins (RBPs) in cancer, with the RBFOX family of RBPs standing out as having the most recurrently down-regulated regulon across multiple cancer types. RBFOX proteins are known regulators of alternative splicing and mRNA stability [22] and have been implicated in a number of neurological diseases [17, 25, 36], but their role in cancer is less characterized. Nonetheless, at least the *RBFOX1* locus appears to be among the most frequently deleted loci across different cancer types [37, 38], with its deletion [39] or specific genetic variations [40] associated with poor survival. Our study suggests that downregulation of RBFOX proteins leads to destabilization of their target transcripts in tumour; many of these transcripts encode proteins involved in calcium signaling, a critical pathway that affects a wide range of cancer-associated processes such as proliferation, invasion, and apoptosis [41]. The association between RBFOX1 and calcium signaling is also supported by previous literature that shows a positive effect of RBFOX1 on the expression of some of the genes involved in this pathway [42].

In addition to RBPs, our results also highlight cancer type-specific deregulation of mRNA stability by miRNAs, with miR-29 standing out as a pan-cancer stability factor. Our observations are in line with previous studies showing that different miR-29 isoforms act as tumor suppressors and are downregulated in several cancer types [43, 44], affecting cell proliferation, differentiation and apoptosis [45]. This downregulation correlates with more aggressive forms of cancer, characterized by increased metastasis, invasion and relapse [46], and therapeutic restoration of miR-29 was suggested to improve disease prognosis [47]. In line with these reports, we observed pan-cancer stabilization of miR-29 targets, suggesting widespread reduction in miR-29 activity in cancer, which could be partially reversed by miR-29 rescue. We note that our results highlight a core set of 64 genes that are bound by miR-29, stabilized at least in KIRC, and down-regulated after restoring miR-29 activity in the KIRC model cell line 786-O (**Fig. 5e**). Importantly, eight of these genes are markers of embryonal carcinoma, suggesting that miR-29 inhibition is essential for activation of an embryonic-like program in cancer. In addition, we observed a significant enrichment of the extracellular matrix (ECM) genes (**Fig. 5e**), suggesting that miR-29 inhibition also contributes to ECM remodeling in cancer, consistent with previous reports on ECM regulation by miR-29 [48].

Together, these results highlight a key role for mRNA stability programs, mediated by RBPs and miRNAs, in regulation of pathways that are integral to cancer development and progression. While the vast majority of current literature is focused on the role of transcriptional mechanisms in reprogramming cancer cells, this study underlines a critical and largely uncharacterized role for post-transcriptional remodeling of the cancer cell transcriptome, and provides a resource for exploring post-transcriptional pathways in cancer.

## Methods

### Joint modelling of intronic and exonic read counts and mRNA stability

Our approach for statistical modeling of intronic and exonic read counts builds on previous research that connects the abundance of pre-mRNA and mature mRNA to mRNA stability (**Fig. S1a-b**):

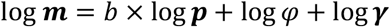

Here, *m* corresponds to the vector of the mature mRNA abundance for a given gene across different samples, *p* is the abundance of the pre-mature mRNA, γ is the mRNA stability across samples, *φ* is the maximum processing rate of RNA, and *b* is the bias-term (**Fig. S1b**). Vectors are differentiated from scalars using bold typeface. We further model the logarithm of mRNA stability as a linear function of a set of sample-level variables:

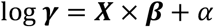

Here,*X* is the *n×k* matrix of sample-level variables (for *n* samples and *k* variables), *β* is the vector of coefficients that quantify the effect of each variable on the mRNA stability, and *α* is an intercept (matrices are differentiated from vectors using capital letters). This leads to:

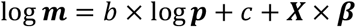

where *c* = log *φ* + *α*. We model the mean of intronic read counts for a given gene across samples as a function of the pre-mRNA abundance for that gene, a gene-level scaling factor that can be interpreted as the effective length, and a sample-specific scaling factor that can be interpreted as library size:

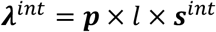

Here, *int* stands for intronic, *λ* represents the mean read count, *l* is the gene-specific scaling factor, and *s* is the sample-specific scaling factor. Similarly, the mean of exonic read counts for a given gene across samples can be expressed as:

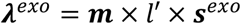

The above equations can be collectively expressed by matrix operations as:

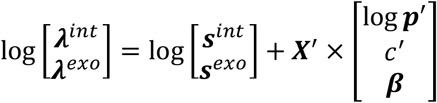

where

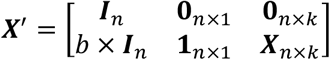

and *p*′= *p* × *l*, *c*′ = *c* + log(*l*′/*l*), and *I* is the identity matrix (matrix dimensions are indicated as subscripts). These equations connect pre-/mature mRNA abundance and mRNA stability to the observed intronic and exonic read counts for each given gene (see **Fig. S1c-d** for matrix equations that consider all genes at the same time). This formulation enables the estimation of unknown parameters using a generalized linear model with a log-link function. In this study, we use DESeq2 [49] to fit the unknown parameters of this model, as explained below.

### Estimation of the effect of sample variables on mRNA stability

The above equations allow us to estimate the distribution of latent variables log *p*′, *c*′, and *β* by fitting the model to observed intronic and exonic read counts. For this purpose, we use the matrix *X*′ as the design matrix in a DESeq2 model. In practice, we replace the first column of *X*′ with an intercept (**Fig. 1c**), which is an equivalent design matrix and does not change the interpretation of *β*, but enables the user to employ a beta prior (if desired) when fitting the DESeq2 model.

In order to be able to construct *X′*, the bias term *b* needs to be first estimated. We do this by first optimizing *b* in order to maximize the likelihood of observed intronic and exonic read counts across all genes in a model that assumes the mRNA stability is a gene-specific constant. Specifically, we use the below design matrix *D* to fit the model using DESeq2, while varying the value of *b* in the interval [0,1] to select the *b* that maximizes the sum of log-likelihood of the data across all genes:

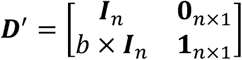

We use the ‘optimize’ function in R to select the optimal value of *b*. Once this optimal value is identified, it is used in the matrix *X*′ (see above), which is then used as the design matrix in DESeq2 to estimate the latent variables, including *β* (i.e. the effect of each variable on stability). This procedure is implemented in DiffRAC (https://github.com/csglab/DiffRAC).

### A modified design to accommodate larger sample sizes

A major limitation of this approach is the considerable increase in computing time with larger sample sizes when DESeq2 is used to fit the model, since the model includes sample-specific latent variables for pre-mRNA abundance. To accommodate these cases, we have also implemented a model that assumes that most of the variance in pre-mRNA abundance can be explained by the experimental variables, instead of including sample-specific latent variables:

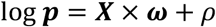

Here, *ω* is the vector of coefficients that represent the effect of each variable on the pre-mRNA abundance of a given gene, and *ρ* is a gene-specific intercept. There, we also have:

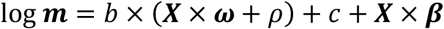

This leads to a modified set of matrix equations (**Fig. S2a-c**) that connect intronic/exonic read counts to sample variables:

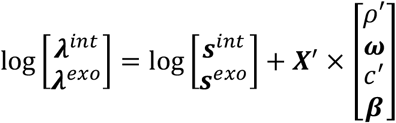

where

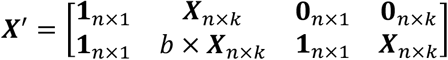

and *ρ*′= *ρ* + log *l*, and *c*′ = *c* + log(*l*′/*l*) + *ρ*×(b–1). Similar to the previous section, *X*′ can be used as the design matrix for DESeq2 to estimate the latent variables, including *ω* and *β*. To construct *X*′, the bia-sterm *b* is chosen so that it maximizes the sum of log-likelihood of data across all genes in a model that assumes gene-specific constant stability, i.e. with the below design matrix *D*′:

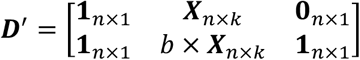

This simplified model is also implemented in DiffRAC. Overall, we see strong agreement between DiffRAC’s estimates when using the two different models (i.e. sample-specific pre-mRNA abundances vs. condition-specific pre-mRNA abundances) on the same data (**Fig. S2d**).

### TCGA RNA-seq data processing

RNA-seq BAM files for 7078 tumor samples and 682 adjacent normal samples from the 18 cancer types with at least 5 normal samples in TCGA were acquired from the National Cancer Institute (NCI) Genomic Data Commons (GDC) data portal (https://portal.gdc.cancer.gov/GDC; dbGaP study accession phs000178.v1.p1). In order to quantify the number of reads corresponding to pre-mRNA and mature mRNA for the estimation of mRNA stability, we generated custom annotations for exons and introns for the transcripts supported by both Ensembl and Havana consortia, using GTF formatted annotations acquired from Ensembl GRCh38 version 87. To avoid the potential confounding effect of alternative splicing on mature mRNA quantification, we exclusively retained exonic reads mapping to constitutive exons that are present in all Ensembl/Havana transcripts. Intronic regions were included in our annotations only if they were not overlapped by any exon, regardless of whether the exon was concordantly annotated by Ensembl or Havana consortia. The strandedness of RNA-seq data was determined using RSeQC [50]. Subsequently, BAM files were sorted by read name using SAMtools, and exonic and intronic reads were separately counted using HTSeq-count [51], limiting to reads with a MAPQ score ≥30. Exonic reads were counted using the HTSeq “intersection-strict” mode, whereas intronic reads were counted using the “union” mode. The exonic/intronic read counts were then used as input to DiffRAC for stability analysis. We removed the cell cycle genes (based on GO term GO:000704) for downstream analyses, given that these genes are not at steady state, which is required for estimating stability from pre-/mature mRNA abundances. When intersecting the DiffRAC KIRC results with miR-29 mimic expression results, however, we kept potential cell cycle genes that were significantly altered by miR-29 mimic expression, as their differential expression is supported by direct measurements in isogenic cell lines.

### Pathway analysis

MSigDB hallmark gene-sets [52] were retrieved using the msigdbr R package (https://cran.r-project.org/web/packages/msigdbr/index.html). For each TCGA cancer type, Fisher’s exact test was used to examine the association between each pathway and the sets of significantly stabilized or destabilized genes, separately.

### Differential RNA stability between MDA-MB-231 and MDA-LM2 cells

Raw RNA-seq reads for time-series measurements of 4-thiouridine (4sU)-labeled RNA [21] from MDA-MB-231 and MDA-LM2 cells were obtained from GEO accession GSE49608 (SRA accession SRP028570). This RNA-seq dataset represents time points 0, 2, 4, and 7h after a 2-hour treatment of cells with 4sU (four replicates for each cell line at each time point). Reads were mapped to the GRCh38 genome assembly using HISAT2 [53], and gene-level read counts for each sample were obtained using HTSeq-count [51] (“intersection-strict” mode) based on Ensembl GRCh38 v87 gene annotations. Differential mRNA stability between the MDA-MB-231 and MDA-LM2 cells were obtained using DESeq2 [49] by modeling the RNA abundances as a function of ~ *c* + *t* + *c*:*t*, where *c* is the cell type (0 for MDA-MB-231 and 1 for MDA-LM2), *t* is the time point, and *c*:*t* is the interaction between cell type and time. In this model, the coefficient of *c* would represent the differential expression between the two cell types (i.e. difference in abundance at time zero); the coefficient of *t* would represent the stability of each gene in the reference cell line MDA-MB-231 (relative to the average of all genes); and the coefficient of the interaction term *c*:*t* would represent the differential stability of each gene between the two cell lines. For each gene, the coefficient of *c*:*t* and associated statistics were retrieved using DESeq2.

### RBP and miRNA regulon analysis

The stability regulons of 35 RBPs (i.e. the set of genes bound and regulated by each RBP) were obtained from a previous publication [22]. The regulons of miRNA families were obtained by identifying exact miRNA seed matches in the mRNA 3’ UTRs. Specifically, 3’ UTR sequences of protein-coding genes were retrieved using the Ensembl GRCh38 version 87 annotations. We limited the analysis to the genes for which a single 3’ UTR, composed of a single exon, was shared across all isoforms, in order to avoid the possible confounding effects of alternative splicing. The miRNA seed sequences (8nt) were retrieved from TargetScan v7.2 [54], limiting to a set of 153 broadly conserved miRNA families (family conservation score ≥1). Exact seed sequence matches in 3’ UTR sequences were identified while limiting the search space to a maximum of 2000 nt downstream of the stop codon.

The regulon enrichment among up-regulated or down-regulated genes was quantified using a logistic regression approach. Specifically, for each cancer type, we modeled the likelihood of being bound by each RBP/miRNA as a function of the gene status, with −1 corresponding to significantly destabilized genes (FDR≤0.05), +1 corresponding to significantly stabilized genes, and 0 corresponding to non-significant genes. To account for the confounding factors that generally affect the number of binding sites of RNA-binding factors (rather than a specific RBP or miRNA; e.g. 3’ UTR length), we used the total number of binding sites of each gene for RBPs or miRNAs as the background. Specifically, we used a generalized linear model of the binomial family, in which the presence of a binding site for the specific RBP or miRNA of interest is considered as “success”, and the presence of binding sites for other RBPs or miRNAs considered as “failures”. These success/failure counts were modeled as a function of the stability status of the gene using the glm function in R.

### Cell culture and transient transfection of miRNA mimics

The established renal cancer cell line 786-O was purchased from the American Type Culture Collection (ATCC; Rockville, MD, USA) and cultured in Dulbecco’s Modified Eagle Medium (DMEM) supplemented with 10% fetal bovine serum (FBS) and 1% penicillin/streptomycin (Life technologies) at 37°C with 5% CO2. Cells (100000 cells/well in 6-well plates) were reverse-transfected in antibiotic-free medium by 25 nM miRNA mimic (ThermoFisher, 4464066) or control mimics (ThermoFisher, 4464058) using Lipofectamine RNAiMAX Reagent (ThermoFisher,13778075) according to the manufacturer’s recommendations.

### RNA isolation and qRT-PCR analysis of miRNAs

Total RNA was extracted using All Prep DNA/RNA/miRNA Universal kit (Qiagen) 48 hours after mimic transfection. LightCycler 480 instrument (Roche) was used to perform qRT-PCR analysis of miR-29 and miR-26 using TaqMan Fast Advanced miRNA Assays (ThermoFisher, 4444557) following guidelines provided by the manufacturer. Expression was reported as Ct values (**Fig. S5**).

### RNA-sequencing and analysis

Library preparation from total RNA was performed using NEB rRNA-depleted (HMR) stranded library preparation kit according to manufacturer’s instructions, and sequenced using Illumina NovaSeq 6000 (100bp paired-end). RNA-seq reads were aligned to the GRCh38 genome assembly using HISAT2 [53], and gene-level read counts were obtained using HTSeq-count [51] (“intersection-strict” mode) based on Ensembl GRCh38 v87 gene annotations. DESeq2 [49] was used to compute differential gene expression between cells expressing the miR-29 mimic and control mimic.

## Declarations

### Ethics approval and consent to participate

Not applicable.

### Consent for publication

Not applicable.

### Availability of data and materials

DiffRAC is available via GitHub at https://github.com/csglab/DiffRAC. Data generated during this study are included in this published article and its supplementary files. Additional data and analysis files are available at http://csg.lab.mcgill.ca/sup/pancancer_stability/ and via Zenodo (doi: 10.5281/zenodo.4404547). RNA-seq data from the miR-29 mimic expression experiment are available via GEO under accession GSE145088. Other data used in this paper are available via their source publications as indicated in the article.

### Authors’ contributions

G.P. and H.S.N. conceived the study, developed the computational methods, analyzed the data, and wrote the manuscript. P.J. and M.R. performed the miRNA-mimic experiment. R.A. contributed to data processing. Y.R. contributed to experimental design and data interpretation. H.S.N. directed the study.

## Acknowledgements

The results published here are in part based on data generated by the TCGA Research Network: https://www.cancer.gov/tcga.

## Description of Supplementary Files

**Supplementary Figures:** This document includes Figures S1-S6, their associated figure legends, and references that are cited within supplementary figure legends.

**Table S1**: Estimation of differential mRNA stability using the RNA-seq data from mouse embryonic stem cells and terminal neurons, along with ground-truth transcript half-life measurements after transcriptional blockage with actinomycin D. DiffRAC estimates are listed in columns 2-7. Ground-truth half-life measurements are in column 8.

**Table S2**: Pan-cancer analysis of differential mRNA stability across TCGA cancers, comparing tumors to normal tissue. DiffRAC estimates for each cancer type are provided in separate sheets.

**Table S3**: Differential stability comparing the MDA-MB-231 and MDA-LM cell lines.

**Table S4**: Clinical data and tumor purity for TCGA samples that were used for analysis of stage- and/or grade-associated stability changes.

**Table S5**: Genes with significant stability changes associated with tumor stage in at least one of the 11 TCGA cancers for which this type of classification was available. Note that no significant associations were identified for READ, and therefore this cancer type has no entries in this table.

**Table S6**: Genes with significant stability changes associated with tumor grade in at least one of the four cancer types for which this type of classification was available.

**Table S7**: Differential gene expression in RBFOX1 knockdown in terminally differentiated neurons. DESeq2 estimates are provided.

**Table S8**: Differential gene expression in the miR-29 mimic relative to control mimic expression in 786-O cells. DESeq2 estimates are provided.

## References

1. Fish L, Navickas A, Culbertson B, Xu Y, Nguyen HCB, Zhang S, Hochman M, Okimoto R, Dill BD, Molina H, et al: Nuclear TARBP2 Drives Oncogenic Dysregulation of RNA Splicing and Decay. Mol Cell 2019, 75:967–981 e969.

2. Fish L, Zhang S, Yu JX, Culbertson B, Zhou AY, Goga A, Goodarzi H: Cancer cells exploit an orphan RNA to drive metastatic progression. Nat Med 2018, 24:1743–1751.

3. Goodarzi H, Liu X, Nguyen HC, Zhang S, Fish L, Tavazoie SF: Endogenous tRNA-Derived Fragments Suppress Breast Cancer Progression via YBX1 Displacement. Cell 2015, 161:790–802.

4. Goodarzi H, Nguyen HCB, Zhang S, Dill BD, Molina H, Tavazoie SF: Modulated Expression of Specific tRNAs Drives Gene Expression and Cancer Progression. Cell 2016, 165:1416–1427.

5. Perron G, Jandaghi P, Solanki S, Safisamghabadi M, Storoz C, Karimzadeh M, Papadakis AI, Arseneault M, Scelo G, Banks RE, et al: A General Framework for Interrogation of mRNA Stability Programs Identifies RNA-Binding Proteins that Govern Cancer Transcriptomes. Cell Rep 2018, 23:1639–1650.

6. Png KJ, Yoshida M, Zhang XH, Shu W, Lee H, Rimner A, Chan TA, Comen E, Andrade VP, Kim SW, et al: MicroRNA-335 inhibits tumor reinitiation and is silenced through genetic and epigenetic mechanisms in human breast cancer. Genes Dev 2011, 25:226–231.

7. Tavazoie SF, Alarcon C, Oskarsson T, Padua D, Wang Q, Bos PD, Gerald WL, Massague J: Endogenous human microRNAs that suppress breast cancer metastasis. Nature 2008, 451:147–152.

8. Vanharanta S, Marney CB, Shu W, Valiente M, Zou Y, Mele A, Darnell RB, Massague J: Loss of the multifunctional RNA-binding protein RBM47 as a source of selectable metastatic traits in breast cancer. Elife 2014, 3.

9. Goodarzi H, Najafabadi HS, Oikonomou P, Greco TM, Fish L, Salavati R, Cristea IM, Tavazoie S: Systematic discovery of structural elements governing stability of mammalian messenger RNAs. Nature 2012, 485:264–268.

10. Yang E, van Nimwegen, E., Zavolan, M., Rajewsky, N., Schroeder, M., Magnasco, M., & Darnell, J. E., Jr: Decay rates of human mRNAs: correlation with functional characteristics and sequence attributes. Genome research 2003, 13:1863–1872.

11. Wada T, Becskei A: Impact of Methods on the Measurement of mRNA Turnover. Int J Mol Sci 2017, 18.

12. Schofield JA, Duffy EE, Kiefer L, Sullivan MC, Simon MD: TimeLapse-seq: adding a temporal dimension to RNA sequencing through nucleoside recoding. Nat Methods 2018, 15:221–225.

13. Blumberg A, Zhao Y, Huang Y-F, Dukler N, Rice EJ, Krumholz K, Danko CG, Siepel A: Characterizing RNA stability genome-wide through combined analysis of PRO-seq and RNA-seq data. 2019.

14. Lugowski A, Nicholson B, Rissland OS: Determining mRNA half-lives on a transcriptome-wide scale. Methods 2018, 137:90–98.

15. Gaidatzis D, Burger L, Stadler MB: Analysis of intronic and exonic reads in RNA-seq data characterizes transcriptional and post-transcriptional regulation. Nat Biotechnol 2015, 33:722–729.

16. La Manno G, Soldatov R, Zeisel A, Braun E, Hochgerner H, Petukhov V, Lidschreiber K, Kastriti ME, Lonnerberg P, Furlan A, et al: RNA velocity of single cells. Nature 2018, 560:494–498.

17. Alkallas R, Fish L, Goodarzi H, Najafabadi HS: Inference of RNA decay rate from transcriptional profiling highlights the regulatory programs of Alzheimer’s disease. Nat Commun 2017, 8:909.

18. Gaidatzis D, Burger L, Florescu M, Stadler MB: Analysis of intronic and exonic reads in RNA-seq data characterizes transcriptional and post-transcriptional regulation. Nat Biotechnol 2015, 33:722–729.

19. Zanzoni A, Spinelli L, Ribeiro DM, Tartaglia GG, Brun C: Post-transcriptional regulatory patterns revealed by protein-RNA interactions. Sci Rep 2019, 9:4302.

20. Joshi A, Van de Peer Y, Michoel T: Structural and functional organization of RNA regulons in the post-transcriptional regulatory network of yeast. Nucleic Acids Res 2011, 39:9108–9117.

21. Goodarzi H, Zhang S, Buss CG, Fish L, Tavazoie S, Tavazoie SF: Metastasis-suppressor transcript destabilization through TARBP2 binding of mRNA hairpins. Nature 2014, 513:256–260.

22. Ray D, Kazan H, Cook KB, Weirauch MT, Najafabadi HS, Li X, Gueroussov S, Albu M, Zheng H, Yang A, et al: A compendium of RNA-binding motifs for decoding gene regulation. Nature 2013, 499:172–177.

23. Jonas S, Izaurralde E: Towards a molecular understanding of microRNA-mediated gene silencing. Nat Rev Genet 2015, 16:421–433.

24. Guo H, Ingolia NT, Weissman JS, Bartel DP: Mammalian microRNAs predominantly act to decrease target mRNA levels. Nature 2010, 466:835–840.

25. Lee JA, Damianov A, Lin CH, Fontes M, Parikshak NN, Anderson ES, Geschwind DH, Black DL, Martin KC: Cytoplasmic Rbfox1 Regulates the Expression of Synaptic and Autism-Related Genes. Neuron 2016, 89:113–128.

26. Weyn-Vanhentenryck SM, Mele A, Yan Q, Sun S, Farny N, Zhang Z, Xue C, Herre M, Silver PA, Zhang MQ, et al: HITS-CLIP and integrative modeling define the Rbfox splicing-regulatory network linked to brain development and autism. Cell Rep 2014, 6:1139–1152.

27. Fogel BL, Wexler E, Wahnich A, Friedrich T, Vijayendran C, Gao F, Parikshak N, Konopka G, Geschwind DH: RBFOX1 regulates both splicing and transcriptional networks in human neuronal development. Hum Mol Genet 2012, 21:4171–4186.

28. Kanehisa M, Goto S: KEGG: kyoto encyclopedia of genes and genomes. Nucleic Acids Res 2000, 28:27–30.

29. Jopling C: Liver-specific microRNA-122: Biogenesis and function. RNA Biol 2012, 9:137–142.

30. Wu C, Zhang J, Cao X, Yang Q, Xia D: Effect of Mir-122 on Human Cholangiocarcinoma Proliferation, Invasion, and Apoptosis Through P53 Expression. Med Sci Monit 2016, 22:2685–2690.

31. Liu N, Jiang F, He TL, Zhang JK, Zhao J, Wang C, Jiang GX, Cao LP, Kang PC, Zhong XY, et al: The Roles of MicroRNA-122 Overexpression in Inhibiting Proliferation and Invasion and Stimulating Apoptosis of Human Cholangiocarcinoma Cells. Sci Rep 2015, 5:16566.

32. Ribatti D, Tamma R, Annese T: Epithelial-Mesenchymal Transition in Cancer: A Historical Overview. Transl Oncol 2020, 13:100773.

33. Meyer N, Penn LZ: Reflecting on 25 years with MYC. Nat Rev Cancer 2008, 8:976–990.

34. Dang CV: MYC on the path to cancer. Cell 2012, 149:22–35.

35. Warburg O, Wind F, Negelein E: The Metabolism of Tumors in the Body. J Gen Physiol 1927, 8:519–530.

36. Lal D, Pernhorst K, Klein KM, Reif P, Tozzi R, Toliat MR, Winterer G, Neubauer B, Nurnberg P, Rosenow F, et al: Extending the phenotypic spectrum of RBFOX1 deletions: Sporadic focal epilepsy. Epilepsia 2015, 56:e129–133.

37. Hu J, Ho AL, Yuan L, Hu B, Hua S, Hwang SS, Zhang J, Hu T, Zheng H, Gan B, et al: From the Cover: Neutralization of terminal differentiation in gliomagenesis. Proc Natl Acad Sci U S A 2013, 110:14520–14527.

38. Rajaram M, Zhang J, Wang T, Li J, Kuscu C, Qi H, Kato M, Grubor V, Weil RJ, Helland A, et al: Two Distinct Categories of Focal Deletions in Cancer Genomes. PLoS One 2013, 8:e66264.

39. Andersen CL, Lamy P, Thorsen K, Kjeldsen E, Wikman F, Villesen P, Oster B, Laurberg S, Orntoft TF: Frequent genomic loss at chr16p13.2 is associated with poor prognosis in colorectal cancer. Int J Cancer 2011, 129:1848–1858.

40. Huang YT, Heist RS, Chirieac LR, Lin X, Skaug V, Zienolddiny S, Haugen A, Wu MC, Wang Z, Su L, et al: Genome-wide analysis of survival in early-stage non-small-cell lung cancer. J Clin Oncol 2009, 27:2660–2667.

41. Monteith GR, Prevarskaya N, Roberts-Thomson SJ: The calcium-cancer signalling nexus. Nat Rev Cancer 2017, 17:367–380.

42. Shen F, Xu X, Yu Z, Li H, Shen H, Li X, Shen M, Chen G: Rbfox-1 contributes to CaMKIIalpha expression and intracerebral hemorrhage-induced secondary brain injury via blocking micro-RNA-124. J Cereb Blood Flow Metab 2020:271678X20916860.

43. He H, Wang L, Zhou W, Zhang Z, Wang L, Xu S, Wang D, Dong J, Tang C, Tang H, et al: MicroRNA Expression Profiling in Clear Cell Renal Cell Carcinoma: Identification and Functional Validation of Key miRNAs. PLoS One 2015, 10:e0125672.

44. Yan B, Guo Q, Fu FJ, Wang Z, Yin Z, Wei YB, Yang JR: The role of miR-29b in cancer: regulation, function, and signaling. Onco Targets Ther 2015, 8:539–548.

45. Park SY, Lee JH, Ha M, Nam JW, Kim VN: miR-29 miRNAs activate p53 by targeting p85 alpha and CDC42. Nat Struct Mol Biol 2009, 16:23–29.

46. Heinzelmann J, Henning B, Sanjmyatav J, Posorski N, Steiner T, Wunderlich H, Gajda MR, Junker K: Specific miRNA signatures are associated with metastasis and poor prognosis in clear cell renal cell carcinoma. World J Urol 2011, 29:367–373.

47. Garzon R, Heaphy CE, Havelange V, Fabbri M, Volinia S, Tsao T, Zanesi N, Kornblau SM, Marcucci G, Calin GA, et al: MicroRNA 29b functions in acute myeloid leukemia. Blood 2009, 114:5331–5341.

48. Sengupta S, den Boon JA, Chen IH, Newton MA, Stanhope SA, Cheng YJ, Chen CJ, Hildesheim A, Sugden B, Ahlquist P: MicroRNA 29c is down-regulated in nasopharyngeal carcinomas, up-regulating mRNAs encoding extracellular matrix proteins. Proc Natl Acad Sci U S A 2008, 105:5874–5878.

49. Love MI, Huber W, Anders S: Moderated estimation of fold change and dispersion for RNA-seq data with DESeq2. Genome Biol 2014, 15:550.

50. Wang L, Wang S, Li W: RSeQC: quality control of RNA-seq experiments. Bioinformatics 2012, 28:2184–2185.

51. Anders S, Pyl PT, Huber W: HTSeq--a Python framework to work with high-throughput sequencing data. Bioinformatics 2015, 31:166–169.

52. Liberzon A, Birger C, Thorvaldsdottir H, Ghandi M, Mesirov JP, Tamayo P: The Molecular Signatures Database (MSigDB) hallmark gene set collection. Cell Syst 2015, 1:417–425.

53. Kim D, Paggi JM, Park C, Bennett C, Salzberg SL: Graph-based genome alignment and genotyping with HISAT2 and HISAT-genotype. Nat Biotechnol 2019, 37:907–915.

54. Agarwal V, Bell GW, Nam JW, Bartel DP: Predicting effective microRNA target sites in mammalian mRNAs. Elife 2015, 4.

55. Subramanian A, Tamayo P, Mootha VK, Mukherjee S, Ebert BL, Gillette MA, Paulovich A, Pomeroy SL, Golub TR, Lander ES, Mesirov JP: Gene set enrichment analysis: a knowledge-based approach for interpreting genome-wide expression profiles. Proc Natl Acad Sci U S A 2005, 102:15545–15550.

56. Pinto JP, Kalathur RK, Oliveira DV, Barata T, Machado RS, Machado S, Pacheco-Leyva I, Duarte I, Futschik ME: StemChecker: a web-based tool to discover and explore stemness signatures in gene sets. Nucleic Acids Res 2015, 43:W72–77.

