## Supplementary Figures for "Pan-cancer analysis of mRNA stability for decoding tumour post-transcriptional programs"

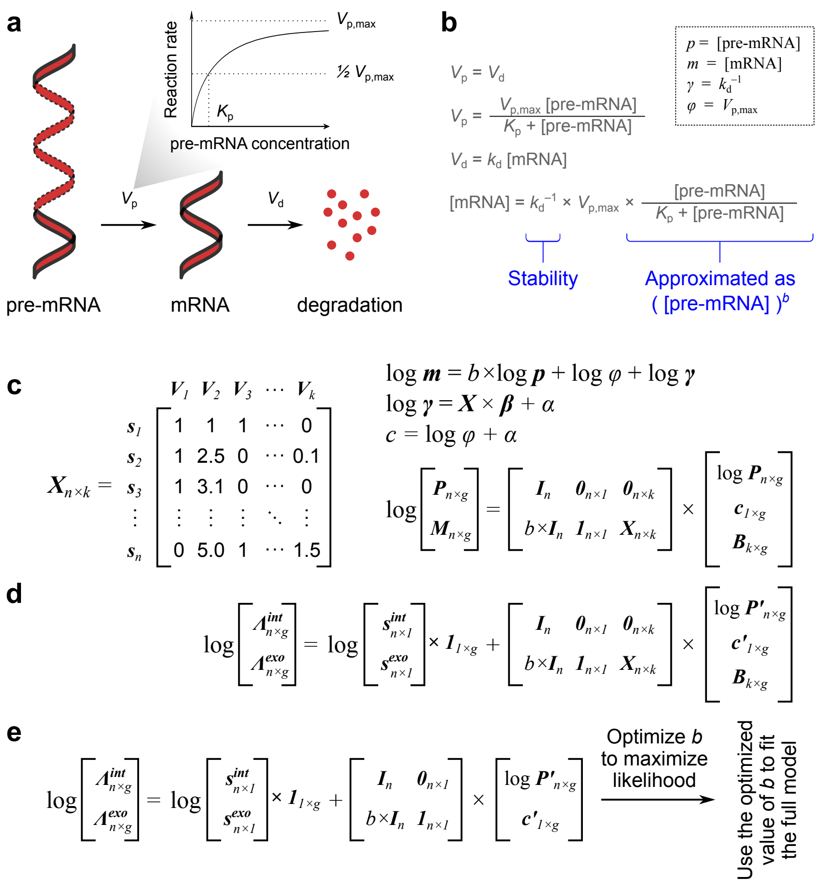


**Figure S1. The DiffRAC model.** (**a**) Schematic presentation of mRNA processing and decay steps based on the model proposed by Alkallas et al. [1]. (**b**) This model leads to an equation that can be approximated as a power-law relationship between stability and the abundance of pre-mRNA and mature mRNAs. Panels **a**-**b** are modified from Alkallas et al. [1] (reproduced under CC-BY license, https://creativecommons.org/licenses/by/4.0/). The notations that are used in this paper to present different quantities in this kinetic model are shown on the top-right. (**c**) DiffRAC aims to model mRNA stability ($\gamma$) as a function of a given a design matrix $X$, which specifies sample characteristics (left). To do this, we start from the power-law relationship between pre-mRNA and mature mRNA abundance and convert it to log-scale (top-right). Logarithm of stability is then modeled as a linear function of the design variables $X$ with coefficients $\beta$. This system of equations can be expressed using matrix operations, as shown on the bottom-right. $X$: the $n\times k$ design matrix for $k$ variables across $n$ samples. $P$: the $n\times g$ matrix of pre-mRNA abundance for $g$ genes across $n$ samples. $M$: the $n\times g$ matrix of mature mRNA abundance. $I_n$: the $n\times n$ identity matrix. $B$: the $k\times g$ matrix of coefficients representing the effect of each of the $k$ variables on the stability of each of the $g$ genes. The parameter $b$ is the bias-term (same as in panel **b**), which is assumed to be shared across genes and samples. The parameter $c$ is a gene-specific factor that represents the combined effect of maximum processing rate ($\varphi$) and baseline RNA stability ($\alpha$, i.e. the intercept for the stability function). (**d**) DiffRAC estimates the latent variables in this model by fitting them to the observed intronic (*int*) and exonic (*exo*) read counts. For this purpose, the logarithm of the mean ($\lambda$) of read counts is modeled as a function of pre-mRNA and mature mRNA abundances (from panel **c**), in addition to sample-specific library size factors ($s$) for intronic and exonic counts. $\Lambda$: the $n\times g$ matrix of the mean of intronic (*int*) or exonic (*exo*) read counts, for $g$ genes across $n$ sampels. Note that here $P$ is replaced with $P'$, as the fitted values will represent the combination of $P$ and a latent, gene-specific scaling factor for intronic reads. Similarly, $c$ is replaced with $c'$ since the fitted values will represent the combination of $c$ and a latent, gene-specific scaling factor for exonic reads. (**e**) To estimate $b$, DiffRAC first fits a model that does not include the effect of stability, and optimizes $b$ to maximize the joint likelihood of intronic and exonic read counts. In other words, $b$ is chosen to maximize the likelihood of data in the absence of changes in mRNA stability. This optimized $b$ is then used to fit the full model in order to examine whether addition of mRNA stability terms to the model significantly improves the fit. DiffRAC uses DESeq2 [2] for library size estimation, model fitting, and likelihood calculation. See **Methods** for more details.

**a**

$$\log \boldsymbol{m} = b\times\log \boldsymbol{p} + \log\varphi + \log\boldsymbol{\gamma}$$

$$\log\boldsymbol{\gamma} = \boldsymbol{X}\times\boldsymbol{\beta} + \alpha \qquad c = \log\varphi + \alpha$$

$$\log\boldsymbol{p} = \boldsymbol{X}\times\boldsymbol{\omega} + \rho$$

$$\log\begin{bmatrix}\boldsymbol{P}_{n\times g}\\ \boldsymbol{M}_{n,g}\end{bmatrix} = \begin{bmatrix}\boldsymbol{1}_{n\times 1} & \boldsymbol{X}_{n\times k} & \boldsymbol{0}_{n\times 1} & \boldsymbol{0}_{n\times k}\\ \boldsymbol{1}_{n\times 1} & b\times\boldsymbol{X}_{n\times k} & \boldsymbol{1}_{n\times 1} & \boldsymbol{X}_{n\times k}\end{bmatrix}\times\begin{bmatrix}\boldsymbol{\rho}_{1\times g}\\ \boldsymbol{\Omega}_{k\times g}\\ \boldsymbol{c}_{1\times g}\\ \boldsymbol{B}_{k\times g}\end{bmatrix}$$

**b**

$$\log\begin{bmatrix}\boldsymbol{\Lambda}^{\boldsymbol{int}}_{n\times g}\\ \boldsymbol{\Lambda}^{\boldsymbol{exo}}_{n\times g}\end{bmatrix} = \log\begin{bmatrix}\boldsymbol{s}^{\boldsymbol{int}}_{n\times 1}\\ \boldsymbol{s}^{\boldsymbol{exo}}_{n\times 1}\end{bmatrix}\times\boldsymbol{1}_{1\times g} + \begin{bmatrix}\boldsymbol{1}_{n\times 1} & \boldsymbol{X}_{n\times k} & \boldsymbol{0}_{n\times 1} & \boldsymbol{0}_{n\times k}\\ \boldsymbol{1}_{n\times 1} & b\times\boldsymbol{X}_{n\times k} & \boldsymbol{1}_{n\times 1} & \boldsymbol{X}_{n\times k}\end{bmatrix}\times\begin{bmatrix}\boldsymbol{\rho}'_{1\times g}\\ \boldsymbol{\Omega}_{k\times g}\\ \boldsymbol{c}'_{1\times g}\\ \boldsymbol{B}_{k\times g}\end{bmatrix}$$

**c**

$$\log\begin{bmatrix}\boldsymbol{\Lambda}^{\boldsymbol{int}}_{n\times g}\\ \boldsymbol{\Lambda}^{\boldsymbol{exo}}_{n\times g}\end{bmatrix} = \log\begin{bmatrix}\boldsymbol{s}^{\boldsymbol{int}}_{n\times 1}\\ \boldsymbol{s}^{\boldsymbol{exo}}_{n\times 1}\end{bmatrix}\times\boldsymbol{1}_{1\times g} + \begin{bmatrix}\boldsymbol{1}_{n\times 1} & \boldsymbol{X}_{n\times k} & \boldsymbol{0}_{n\times 1}\\ \boldsymbol{1}_{n\times 1} & b\times\boldsymbol{X}_{n\times k} & \boldsymbol{1}_{n\times 1}\end{bmatrix}\times\begin{bmatrix}\boldsymbol{\rho}'_{1\times g}\\ \boldsymbol{\Omega}_{k\times g}\\ \boldsymbol{c}'_{1\times g}\end{bmatrix}$$

Optimize *b* to maximize likelihood → Use the optimized value of *b* to fit the full model

**d**

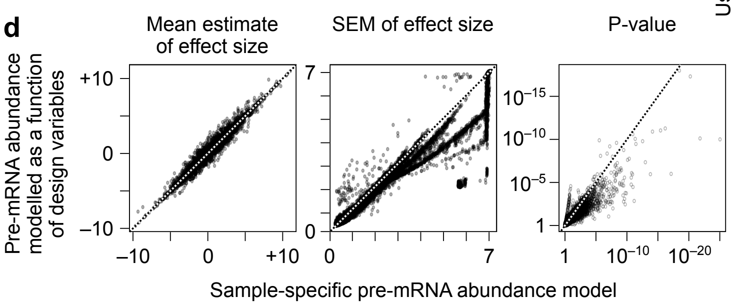


**Figure S2. A simplified version of the DiffRAC model.** (**a**) In this simplified model, instead of assuming sample-specific latent pre-mRNA abundances, the pre-mRNA abundance is also modeled as a linear function of the design matrix *X*, with the coefficients $\omega$ and intercept (baseline pre-mRNA abundance) equal to $\rho$. $\Omega$: the $k\times g$ matrix of coefficients representing the effect of each of the *k* variables on the pre-mRNA abundance of each of the *g* genes. Other variables are the same as **Fig. S1**. (**b**) Similar to **Fig. S1d**, DiffRAC fits this model to the observed intronic and exonic read counts. Note that $\rho$ and *c* are replaced with $\rho'$ and $c'$ since the fitted values will also contain latent gene-specific scaling factors for intronic and exonic counts, respectively. (**c**) Similar to **Fig. S1e**, the bias parameter *b* is estimated by maximizing the likelihood for a model that assumes constant mRNA stability for each gene across samples. This optimized *b* is then used in the full model. (**d**) DiffRAC produces comparable statistics when sample-specific latent variables are included for pre-mRNA abundance (i.e. the full model; x-axis) or pre-mRNA abundance is modeled as a function of design variables (i.e. the simplified model; y-axis). Data correspond to DiffRAC analysis of differential stability between mouse ES cells and ES cells differentiated to terminal neurons [3].

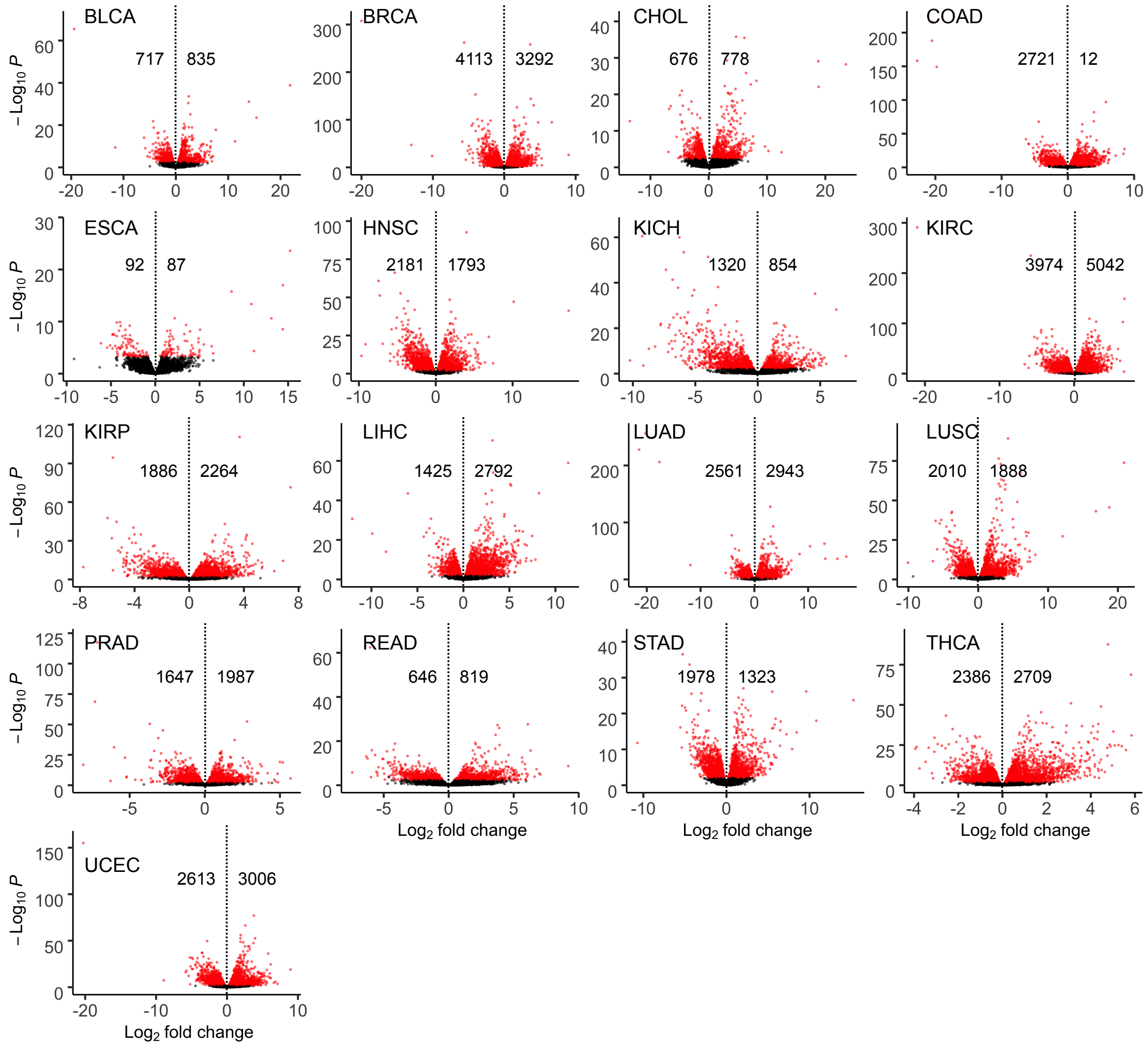


**Figure S3. Differential mRNA stability in TCGA cancers**. Volcano plots showing differential stability, between tumour and normal samples, in each of the 18 TCGA cancers analyzed by DiffRAC. The x-axis represents the $\log_2$ fold-change of mRNA stability (T/N) and the y-axis represents the $\log_{10}$ P. Genes that pass FDR < 0.05 are colored in red. The number of significantly stabilized (right) and destabilized (left) genes are indicated on the plots.

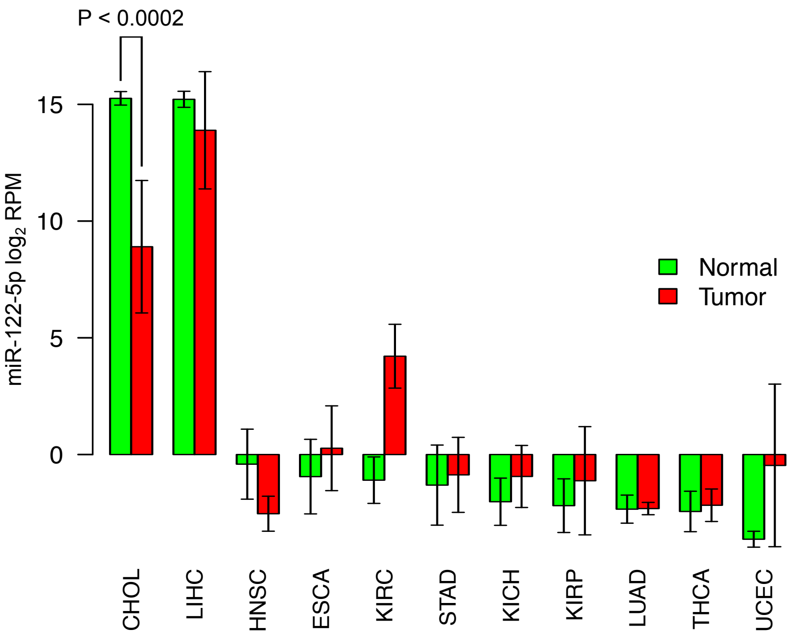


**Figure S4. Expression of miR-122 across cancer types**. Only TCGA cancer types with miR-122 measurement in patient-matched normal and tumour tissues are included. P-value is based on paired t-test.

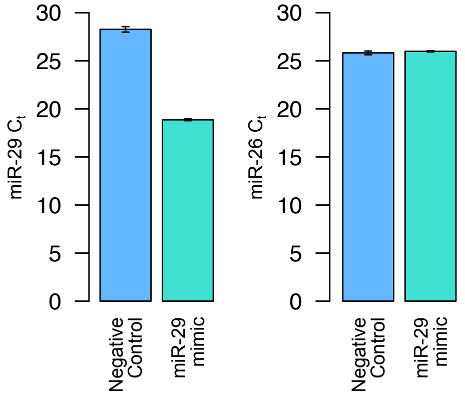


**Figure S5. qRT-PCR confirmation of miR-29-mimic expression in 786-O cells**. Negative control corresponds to transfection with a control mimic. Left: change in miR-29b expression after transfection with miR-29b-mimic. Right: change in miR-26 expression (as an internal control). Average $C_t$ values (n=3) are shown, with the error bars corresponding to the standard deviation. Note that smaller $C_t$ values correspond to higher expression of the assayed miRNA.

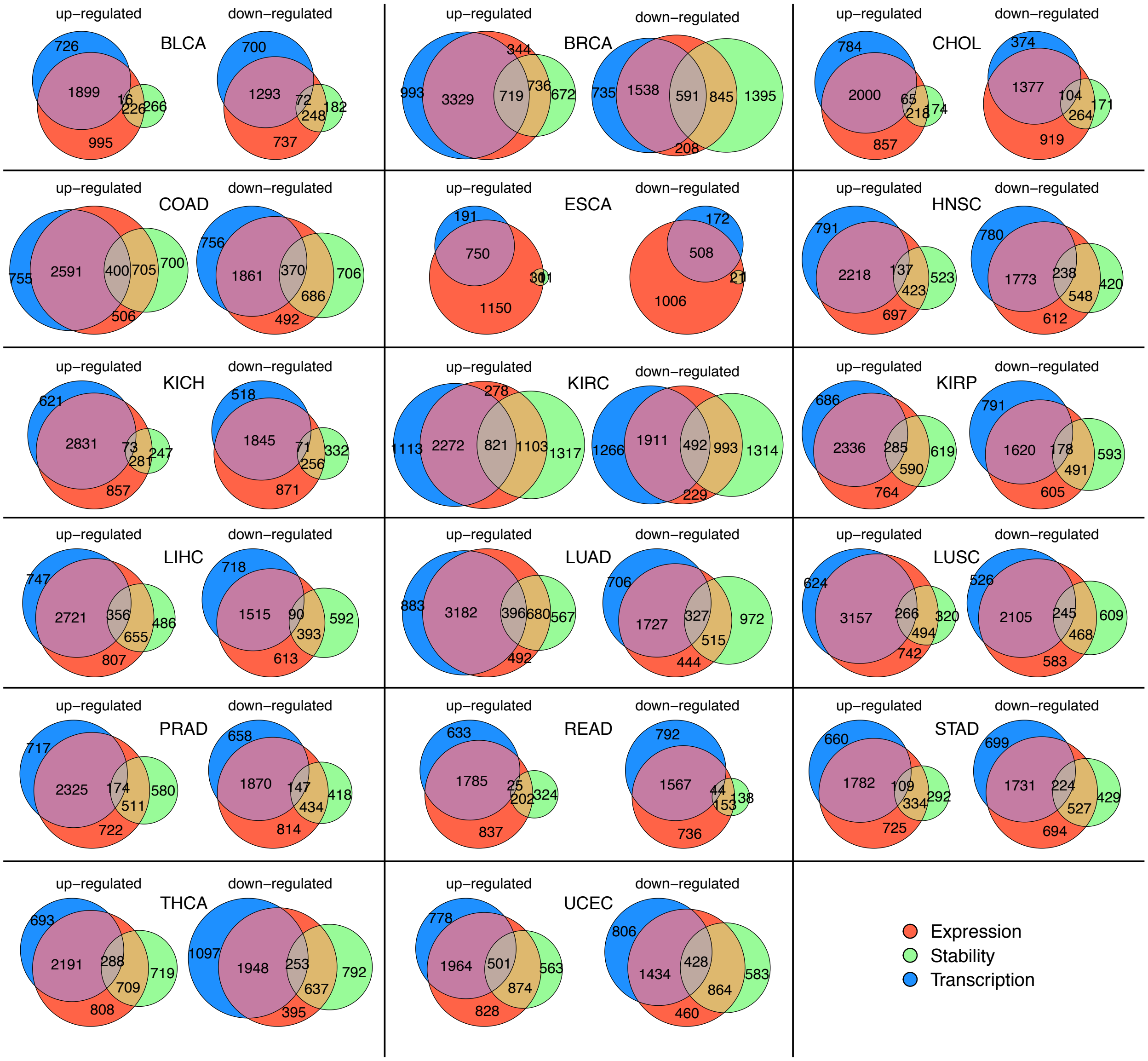


**Figure S6.** Overlap of differentially expressed genes with genes that show differential stability or differential transcription in each cancer. For each cancer type, differentially up-regulated or down-regulated genes are plotted separately. In each case, differential expression was assayed using DESeq2 based on read counts downloaded from TCGA. Differential stability is based on DiffRAC. Differential transcription is based on DiffRAC variables that correspond to pre-mRNA abundance. In all cases, significant hits are identified based on FDR < 0.05.

1. Alkallas R, Fish L, Goodarzi H, Najafabadi HS: **Inference of RNA decay rate from transcriptional profiling highlights the regulatory programs of Alzheimer's disease.** *Nat Commun* 2017, **8:**909.
2. Love MI, Huber W, Anders S: **Moderated estimation of fold change and dispersion for RNA-seq data with DESeq2.** *Genome Biol* 2014, **15:**550.
3. Gaidatzis D, Burger L, Florescu M, Stadler MB: **Analysis of intronic and exonic reads in RNA-seq data characterizes transcriptional and post-transcriptional regulation.** *Nat Biotechnol* 2015, **33:**722-729.
